# Highly accurate random DNA sequencing using inherent interlayer potential traps of bilayer MoS_2_ nanopores

**DOI:** 10.1101/2020.04.21.053595

**Authors:** Payel Sen, Hiofan Hoi, Dipanjan Nandi, Manisha Gupta

## Abstract

Solid-state MoS_2_ nanopores are emerging as potential real-time DNA sequencers due to their ultra-thinness and pore stability. One of the major challenges in determining random nucleotide sequence (unlike polynucleotide strands) is the non-homogeneity of the charge interaction and velocity during DNA translocation. This results in varying blockade current for the same nucleotide, reducing the sequencing confidence. In this work, we studied the inherent impedance-tunability (due to vertical interlayer potential gradient and ion accumulation) of multilayered MoS_2_ nanopores along with its effect on improving analyte capture and charge interaction, for more sensitive and confident sensing. Experimentally we demonstrate that 2-3 nm diameter bilayer MoS_2_ pores are best suited for high accuracy (~90%) sequencing of mixed nucleotides with signal-to-noise-ratio greater than 11 in picomolar concentration solutions. High temporal resolution demonstrated by bilayer MoS_2_ nanopores can help detect neutral proteins in future. The high accuracy detection in low concentration analyte can hence be applied for control and prevention of hereditary diseases and understanding health effects of rare microbial strains.

## Introduction

Accurate DNA sequencing can determine genetic susceptibility to hereditary diseases and determine health effects of a microbial strain. Nanopore sequencing introduced in late 1900s was one of the key breakthrough technologies for DNA sequencing due to its real-time, low-cost, long-read, label-free and amplification-free sensing approach^1^. Biological nanopores have been extensively used for sequencing because they facilitate slow DNA translocation, favourable for highly resolved sensing. However, they lack longevity and thermomechanical stability^2^, leading to the development of solid-state nanopores. Solid-state nanopores are commonly fabricated on insulating membranes like silicon nitride (SiN_x_) or silicon dioxide (SiO_2_) by transmission electron microscopy or helium ion microscopy^3–7^. Solid-state nanopores offer the advantage of being stable even though they suffer from high velocity of DNA translocation^2^. In sequencing, analyte DNA molecules are electrophoretically driven through the nanopore and are electrically detected via the ionic current change caused during their passage through the nanopore. Particularly in DNA, each nucleotide (nt) base (adenine, thymine, cytosine, and guanine) can produce distinguishable ionic blockade currents because of the difference in their size and surface charge^6–9^. The characteristic current drops obtained can therefore identify the order of nucleotides, which can be used towards determining the whole DNA sequence^3^. Nanopore sequencing method is often prone to missing individual or few nucleotides leading to erroneous sequence. Sensitivity and the sensing accuracy depend upon the analyte concentration which influences the capture rate, the translocation velocity which decides the residence time in the pore and the nanopore geometry and material which governs the pore conductance. Hence, designing a pore and the measurement setup are very important for measurement of low concentration analyte detection with high accuracy and sensitivity.

Using nanopore technology, DNA can be sequenced with improved accuracy without the need for amplification techniques like Polymerase Chain Reaction (PCR)^10^. However, the probability of DNA occurring in the nanopore capture volume reduces with low sample concentration and pore volume. Achieving a good detection rate and accuracy at picomolar DNA concentrations thus requires optimized surface charge interaction around the nanopore. Few works have been performed to improve sensing efficiency at low concentration^11–13^. Pathogen identification and DNA oligo sensing by Surface Enhanced Raman Spectroscopy has been performed^11–12^. Low concentration analyte detection has been achieved by employing additional techniques like dielectrophoretic (DEP) trapping to increase the electrophoretic pull exerted on the molecular strand to be sensed^13^. The traps help in regulating nanopore volume in a range comparable to the translocating molecules and also increases the capture rate. DEP trapping has been used to sense single nucleotides at a concentration as low as 5 fM^13^. However, these techniques require additional process aids making it more complex. Usually DNA concentrations in micromolar to nanomolar range are employed for solid-state nanopore sequencing^14–19^. In this work we aim at introducing a simple and repeatable approach towards picomolar concentration DNA sequencing by exploiting properties of a suitable membrane material. Thus, one can conduct DNA profiling of dilute samples collected from crime scenes (50 ng/µL to 0.023 ng/µL)^20^, archaeological sites (4-17 ng/ µL)^21^ and even for rare pathogen strain analysis^22–23^, using this technique.

A single dominant mutated gene in each cell can cause harmful health effects. Diseases like cystic fibrosis, sickle-cell anaemia, neurofibromatosis, color-blindness are all caused by point mutations to the genome^24–25^. Identification of single nucleotide mutation can be done if accurate DNA sequence of the mutated gene can be obtained. The mutated sequence can then be compared with wild gene to identify the mutation. This identification of mutated gene can thereby help in early detection of life-threatening diseases. Sensing (blockade current) resolution can be along the space (obtaining current blockades specific to individual bases) and time regime (obtaining high dwell time of each base inside the nanopore)^9, 24^. The current blockade depends on the fraction of the nanopore volume occupied by the analyte at any time instant. In other words, the nanopore volume should be comparable to single nucleotide size to sense individual species. Single nucleotide is 1.6 - 1.8 nm long and ~1 nm in diameter^26^. Therefore, 2 - 4 nm diameter nanopores on ~1.5 - 2 nm thin membrane should show good nucleotide sensing resolution. Silicon nitride (SiN_x_) is one of the most studied nanopore materials till date, especially due to stability and compatibility with standard fabrication techniques^27^. However, higher thickness of the membranes (20-50 nm) compared to molecular dimensions (~1 nm), limits spatial resolution. SiN_x_ membranes thinner than 5 nm have more chances of pits in them, making it difficult to reduce the thickness of the membranes beyond this^28^. In two-dimensional (2D) materials the layers are separated by weak van der Waals forces, which help obtain stable, monolayer thickness control. 2D material nanopores, having a small control volume can therefore be utilized for developing nanopores with geometry desired for single nucleotide detection^29^. Nanopores on molybdenum disulphide (MoS_2_), a type of 2D metal dichalcogenide, have proved to be promising DNA sequencers for their structural stability in aqueous medium, high signal-to-noise ratio and no sticking of DNA^14^. The abundance of Molybdenum (Mo) around MoS_2_ pore offers favorable analyte/pore wall surface charge interactions and minimizes noise, making it a good alternative for nanopore fabrication^14, 30–32^.

There are limited number of studies on single nucleotide sensing using 1-layer thick MoS_2_ nanopores^33–37^. Single nucleotide detection from homogenous polynucleotide strands has been demonstrated by 1-layer thick MoS_2_ nanopores at a concentration of 5 µg ml^−1^ with 100 mM KCl solution^14^. In this work, we have demonstrated single nucleotide sequencing from DNA template containing customized random nucleotide sequence, which demands slower translocation for enhanced surface charge interaction along nucleotide/pore interface. 1-layer thick MoS_2_ makes DNA translocation very fast hampering temporal resolution. One of the major reasons for this is the thickness of 1-layer MoS_2_ is 0.65 nm, which is less than single nucleotide size (1.6-1.8 nm). This compromises charge interaction making DNA translocation fast, thus limiting temporal resolution. Research has been carried out to reduce speed of translocation by introducing viscosity gradient between cis and trans electrolytic chambers^38–42^. In this work, we explore the inherent ability of multilayer (2 - 6 layers thick) MoS_2_ having thickness in the range of 1 - 4 nm, in slowing DNA translocation. Additionally, the behavior of 1-layer and multilayer thick MoS_2_ differs under a vertical electrical field imposed by two electrodes dipped in cis and trans chambers. In multilayer MoS_2_, the strong interlayer coupling creates a potential gradation along the van der Waals separated layers, as each layer experiences a different electric potential^43^. This potential gradient causes trapping and detrapping of the phosphate groups of DNA strand with the Molybdenum atoms at the nanopore. The increased negative-positive charge interaction creates an additional electrophoretic pull^43–44^. This can provide a platform for effective capture and detection of molecules at sub-nanomolar molecular concentrations without using any external trapping mechanism. Thus, by tuning the number of layers of MoS_2_ we can control the velocity and the conductance of the pore.

This work primarily focuses on determining MoS_2_ nanopore properties suitable for label-free sensing of nucleotides from a DNA template containing mixed nucleotides at low (picomolar) concentration with a good detection rate and resolution. Picomolar concentration used for sensing exploits the amplification-free aspect of nanopore sensing enabling high-accuracy, low-cost rare DNA fingerprinting. COMSOL Multiphysics 5.4 simulation followed by experimental recordings were performed for a stepwise optimization of number of layers suitable for a highly repeatable and efficient DNA nucleotide sensing. Here, we have demonstrated both via simulations and experiments that the 2-layers thick MoS_2_ nanopores are better suited for improving detection sensitivity and accuracy for low (picomolar) concentration analytes.

## Results and discussions

### Optimization of the number of MoS_2_ layers for single nucleotide detection

DNA translocation through 1 - 10 nm diameter pores was first simulated using COMSOL Multiphysics 5.4, for different membrane types: 1 - 6 layers MoS_2_ (0.65 - 4 nm thick), 50 nm SiN_x_ and 76 - layers MoS_2_ having thickness (49.4 nm) similar to that of SiN_x_ membrane. A 2D symmetric geometry representing two electrolyte (300 mM KCl) filled half-cells separated by a pore bearing membrane was considered, similar to the experimental setup used as shown in **Figure S5** in Supplementary information. DNA was modelled as a 10 nm long homogenous rectangular poly-nucleotide strand and was introduced in the cis chamber with a trans bias ranging from −200 to 200 mV to pull the DNA through the nanopore. Figure 1.a provides a schematic representation of the force balance conducted on the DNA for simulating blockade conductance and translocation kinetics. Detailed information on modelling geometry (**Figure S1 of Supplementary information**), meshing parameters and physics are discussed in Supplementary information. **Figure S2** shows the 3D revolved geometry with the potential gradient profile for different number of MoS_2_ layers and 50 nm thick SiN_x_ pores.

**Figure 1.**
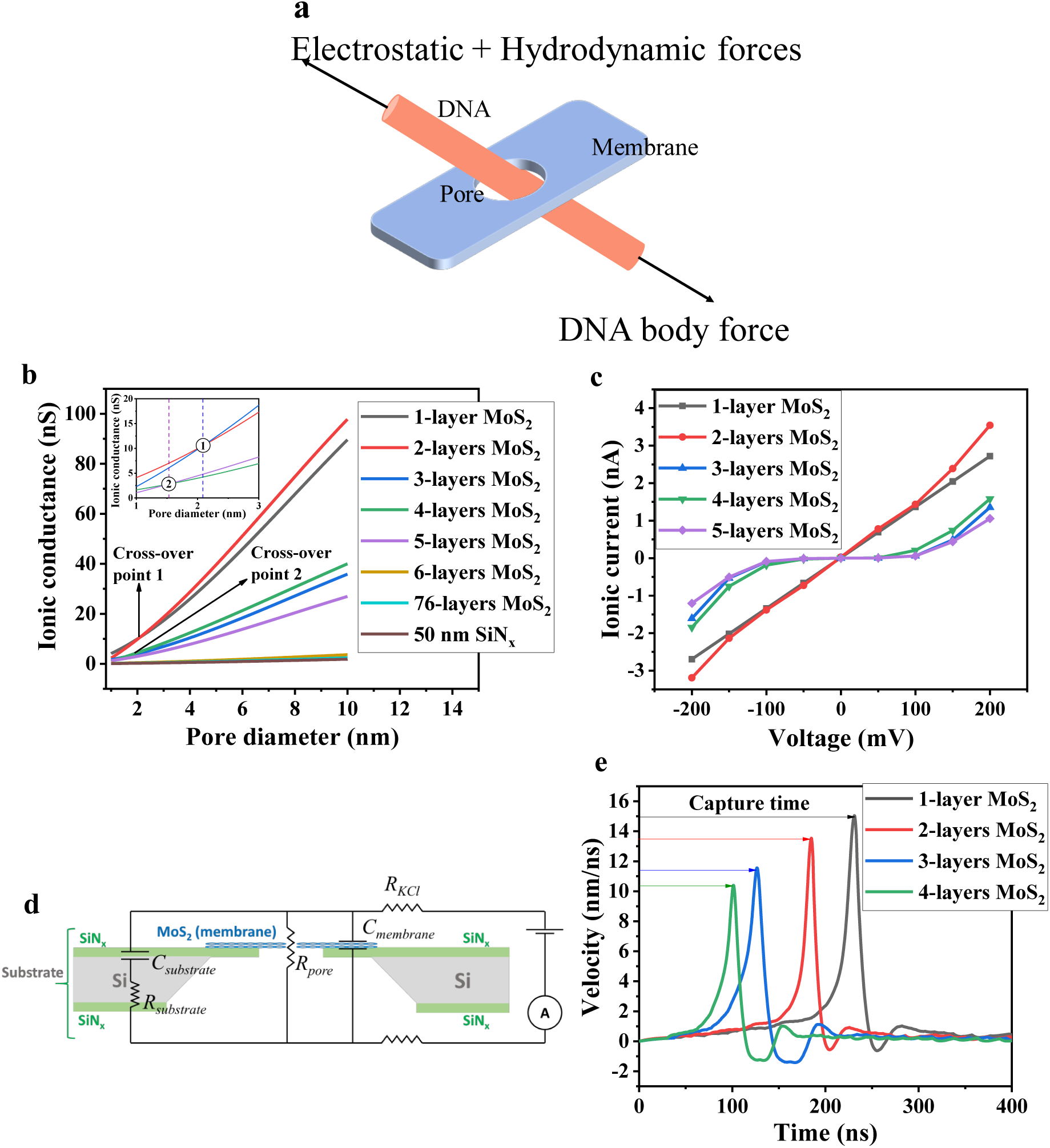
**a.** Simple schematic representation of the force balance used for simulating DNA translocation electrokinetics through nanopores, **b.** Simulated ionic conductance of 1-6 layers MoS_2_ (thickness range: 0.65-4 nm), 50 nm thick SiN_x_ and 76-layers MoS_2_ (thickness: 49.4 nm), showing cross-over points, Cross-over 1 indicates the diameter beyond which ionic conductance for 2-layers thick MoS_2_ nanopore is higher than 1-layer thick MoS_2_. Similarly, above cross-over point 2, the ionic conductance through 4 - layers thick MoS_2_ pore is found to be greater than that for 3-layers thick MoS_2_; inset showing the cross-over points clearly: data reported for −200 mV to 200 mV trans bias and 300 mM KCl **c.** Simulated I-V profile for 1- 4 layers thick MoS_2_ (shortlisted from ionic conductance comparison) for 2.5 nm pore diameter (greater than cross-over point 1), showing highest non-linearly varying I-V response demonstrated by 2-layers thick MoS_2_: simulation conducted for −200 mV to 200 mV bias range and 300 mM KCl, **d.** Representative nanopore circuit showing the active capacitance and resistances, **e.** Simulated translocation velocity profile of 10 nm poly-guanine strand through 1-4 layers thick MoS_2_ nanopores of 2.5 nm diameter, showing the capture time and velocity profile for each pore: data reported for 200 mV bias and 100 mM KCl.

The ionic conductance of these solid-state pores is first simulated for an electrolyte (KCl) concentration of 300 mM and −200 mV to 200 mV trans bias range, while keeping the cis chamber grounded. Figure 1.b shows the variation of ionic conductance with pore diameter for different types of membranes. The cross-over diameter mentioned in Figure 1.b is found to decrease with increase in number of MoS_2_ layers. Cross-over point 1 occurs between 1-layer and 2-layers thick MoS_2_ at 2 nm pore diameter. This necessarily states that ionic conductance better for a 2 – layers thick MoS_2_ pore as compared to the 1-layer thick MoS_2_ pore for diameters greater than 2 nm. Table 1 quantitatively lists the simulated ionic conductance for different thickness MoS_2_ and 50 nm thick SiN_x_ nanopores, with the corresponding I-V displayed in Figure 1.b.

**Table 1:** Simulation results for drawing an ionic conductance-based comparison between different type of nanopores (1 - 6 layers thick MoS_2_, 76 - layers thick MoS_2_ and 50 nm thick SiN_x_) for 200 mV bias and 300 mM KCl. The ionic conductance for different nanopore types having 2.5 nm diameter are listed. The diameters at cross-over points 1 and 2 (extracted from Figure 1.b) are also numerically mentioned below.

| Nanopore type | Membrane thickness | Cross-over points | Ionic conductance |
| --- | --- | --- | --- |
| 1-layer thick MoS<sub>2</sub> | 0.65 nm | Point 1: 2 nm diameter | 13.48 nS |
| 2-layers thick MoS<sub>2</sub> | 1.3 nm |  | 14.12 nS |
| 3-layers thick MoS<sub>2</sub> | 1.9 nm | Point 2: 1.5 nm diameter | 5.39 nS |
| 4-layers thick MoS<sub>2</sub> | 2.6 nm |  | 6.33 nS |
| 5-layers thick MoS<sub>2</sub> | 3.8 nm | — | 4.05 nS |
| 6-layers thick MoS<sub>2</sub> | 5.2 nm |  | 0.57 nS |
| 76-layers thick MoS<sub>2</sub> | 49.4 nm | — | 0.37 nS |
| 50 nm thick SiN<sub>x</sub> | 50 nm | — | 0.28 nS |

The I-V behaviour for the selected 1- 4 layers thick MoS_2_ pores with applied bias from −200 mV to 200 mV in 300 mM KCl was then simulated. 1-layer thick MoS_2_ pores demonstrate a linear I-V curve as compared to the 2 - 4 layers thick MoS_2_ pores show non-linear behaviour (see Figure 1.c). It was found that I-V profile becomes more non-linear with increasing number of layers. The ionic conductance trend and I-V response (Figure 1.b-c) observed by COMSOL Multiphysics simulation can be best explained by utilising a Resistance (R) – Capacitance (C) equivalent model for the nanopore (Figure 1.d). The nanopore acts as a resistance (R) and membrane along with the electrolyte is the capacitance (C). The current variation with respect to resultant impedance (Z) can therefore be studied (see Equation 1 ^[45]^).

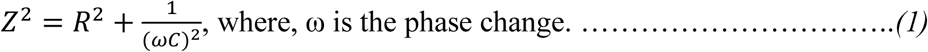

The Cross-over points 1 and 2 indicate the diameters above which the capacitance is dominant as compared to the pore resistance, which reduces the impedance, thus causing a rise in ionic conductance. It is also important to note that this behaviour is highly dependent on the applied bias (greater than 50 mV in this case as seen in Figure 1.c). In case of 1-layer thick MoS_2_, the pore resistance dominates the impedance producing a linear I-V (Figure 1.c). For smaller pores (~1-2 nm diameter), due to higher pore resistance, the impedance increases, thus reducing the ionic current (Figure 1.c). With increase in the number of layers, an interlayer potential gradient along the MoS_2_ interfaces causes accumulation of ions therefore producing a capacitance ^[45–46]^. Hence, the resultant impedance has a contribution from the membrane-based interlayer and intralayer charge storage, electro-activity caused by pseudo-capacitance and the pore resistance ^[45–46]^. As a result, an increase in I-V is observed above a certain applied voltage indicating an increased conductance (Figure 1.c).

To understand sensing performance of SiN_x_ and multilayer MoS_2_ nanopores, the kinetics for translocation of the same analyte DNA were then simulated using 100 mM KCl electrolyte. The results show that increase in number of MoS_2_ layers shortens nucleotide capture time and improves dwell time of the nucleotides at the nanopore (as depicted by lower peak velocity). Figure 1.e shows the translocation velocity profile of homogenous poly-guanine strand through 1-4 layers thick MoS_2_ nanopores. Increased capture rate and longer pore dwell times suggest an improved event rate, sensitivity and better temporal resolution (slower translocation). Higher capture rate and dwell times displayed by 2 and 4-layers thick MoS_2_ pores can therefore be directly related to their sensing efficiency. **Figure S3** and **Figure S4** of Supplementary information respectively shows the conductance blockades and translocation velocity profiles of different poly-nucleotide strands.

### Experimental considerations based on simulated optimizations

To verify our simulation results, we fabricated multiple MoS_2_ nanopores with different number of layers, the procedure of fabrication is explained in detail in the experimental details section. Since, the highest crossover pore diameter observed in ionic conductance simulation for 1-4 layers MoS_2_ (Table 1) was above 2 nm, pores of diameter around 2.5 nm were fabricated by Scanning Transmission Electron Microscopy (STEM). Based on the simulation results, we decided to fabricate and experimentally study 2-layers and 4-layers thick MoS_2_ pores to utilize their higher ionic conductance (as seen in Figure 1.b-c). Experiments were conducted with these pores using low (~ 40 pM) concentration of analyte DNA solution to verify the improved sensitivity (Figure 1.e),

### MoS_2_ membrane and nanopore fabrication and characterization

Free-standing mechanically exfoliated MoS_2_ membranes on SiN_x_ support were characterized by Raman Spectroscopy and High-Resolution Transmission Electron Microscopy (HRTEM) to determine the number of layers of MoS_2_. Nanopores fabricated on 2- and 4-layer thick MoS_2_ were then used for all the other experiments. STEM diffractogram is very accurate for detection of the number of MoS_2_ layers but cannot be used for more than 3 layers. Hence, Raman shift was used to determine 4 – layers thick MoS_2_. Figure 2.a shows the STEM image and diffractogram of a typical exfoliated free-standing 2-layers thick MoS_2_ membrane. Figure 2.b represents Raman spectrum of a typical 4-layer thick MoS_2_ membrane showing a Raman shift of 23.9 cm^−1^, which confirms the number of layers.

**Figure 2.**
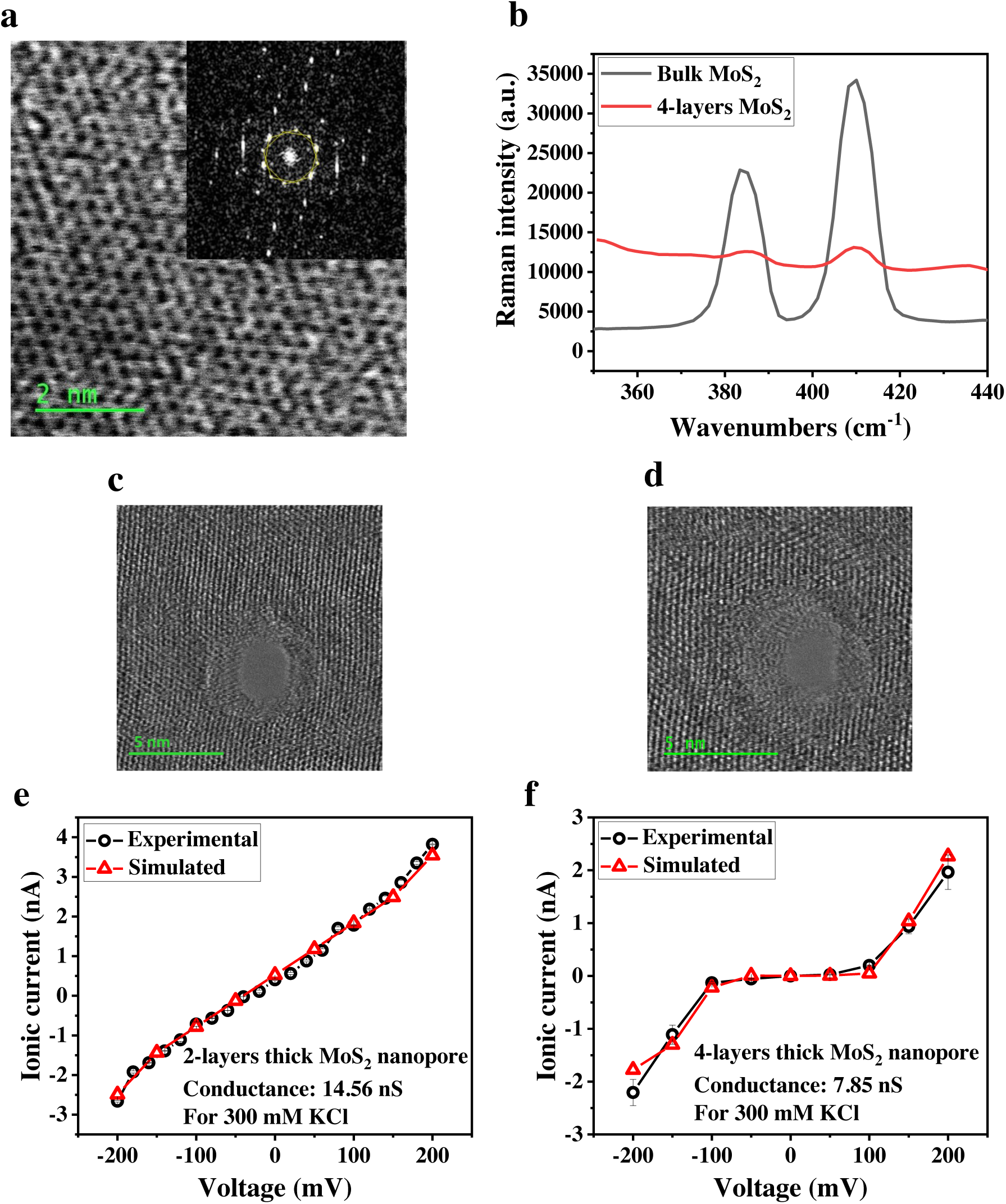
**a.** HRTEM characterization of 2-layers thick MoS_2_ membrane with inset diffractogram showing two overlapped hexagons confirming 2 - layers MoS_2_, **b.** Raman Spectroscopic profile of a representative exfoliated 4 - layers thick MoS_2_ membrane shows two vibrational modes: intra-layer (mainly chemical composition dependent) and inter-layer (large influence by mass and hence number of layers); two peaks at 383 cm^−1^ and 408 cm^−1^ are fingerprints of bulk MoS_2_; the peak distance of 23.9 cm-1 indicates that it consists of 4 layers, **c.** TEM images of STEM fabricated nanopores on 2-layers thick MoS_2_ and **d.** TEM images of STEM fabricated nanopores on 4-layers thick MoS_2_ membranes, **e.** Experimental and simulated I-V profiles (using 300 mM KCl) through 2-layers thick MoS_2_ predicting a 2.5 nm pore diameter and **f.** Experimental and simulated I-V profiles (using 300 mM KCl) through 4-layers thick MoS_2_ pores predicting a 2.8 nm pore diameter by experimental and simulated pore conductance comparison; error bars in **e-f** represents error between multiple conductance experiments.

Nanopores were fabricated using STEM on 2 and 4-layers thick MoS_2_, HRTEM images of which are shown in Figure 2.c-d respectively. Ionic current through the fabricated nanopores was measured by sweeping the voltage from −200 mV to 200 mV and their conductance was then extracted. Each pore was measured 8 times to obtain a stable conductance after repeated cleaning between every measurement, and the average conductance starting from the third measurement was reported. Conductance of 2 and 4-layers thick MoS_2_ nanopore were measured to be ~14.56 nS and ~7.85 nS respectively. We used our simulation results for this conductance to predict the pore diameters as 2.5 nm and 2.8 nm for the 2 and 4 – layers thick MoS_2_ pores respectively. Figure 2.e-f shows the fitted experimental and simulated I-V graphs for the predicted pore diameters.

### DNA sequencing at picomolar concentration (pM) using 2-layers and 4-layers MoS_2_ nanopores

Once the pores conductance was characterised, DNA translocation experiments were conducted on the same pores. For the DNA translocation experiments, we have used a 40 pM solution of single-stranded (ss) DNA oligos containing 30 mixed nucleotides as described in the experimental details section. A bias of 200 mV was used to measure the ionic current of DNA translocation through the 7 fabricated pores each, for 2 and 4-layers thick MoS_2_ having diameters in the range of 2.5 - 3 nm. The measurement results were then used to determine the efficiency and accuracy of the order and count of nucleotides present in the DNA strand.

The recorded translocation profile was analysed to obtain current drop and dwell times. Figure 3.a shows distinct single nucleotide peaks obtained by sequencing through 2-layers thick MoS_2_ nanopores. Figure 3.b shows DNA translocation peaks through 4-layers thick MoS_2_ nanopores showing simultaneous detection of multiple nucleotides. Figure 3.c-d show 4-layers thick MoS_2_ demonstrating higher dwell times and lower current drops compared to 2-layers thick MoS_2_ pores, which agrees with our simulation results. Figure 3.c-d also shows the ability of 2-layers thick MoS_2_ pore for detection of single-nucleotide with well-defined and distinct current levels for different nucleotides whereas the 4-layers thick MoS_2_ pore demonstrates overlapped current signatures for different nucleotides. Thus, it can clearly be seen that the number of layers of MoS_2_ very strongly decide the sensitivity of a pore with 2-layers thick pores showing the best results in terms of single nucleotide detection.

**Figure 3.**
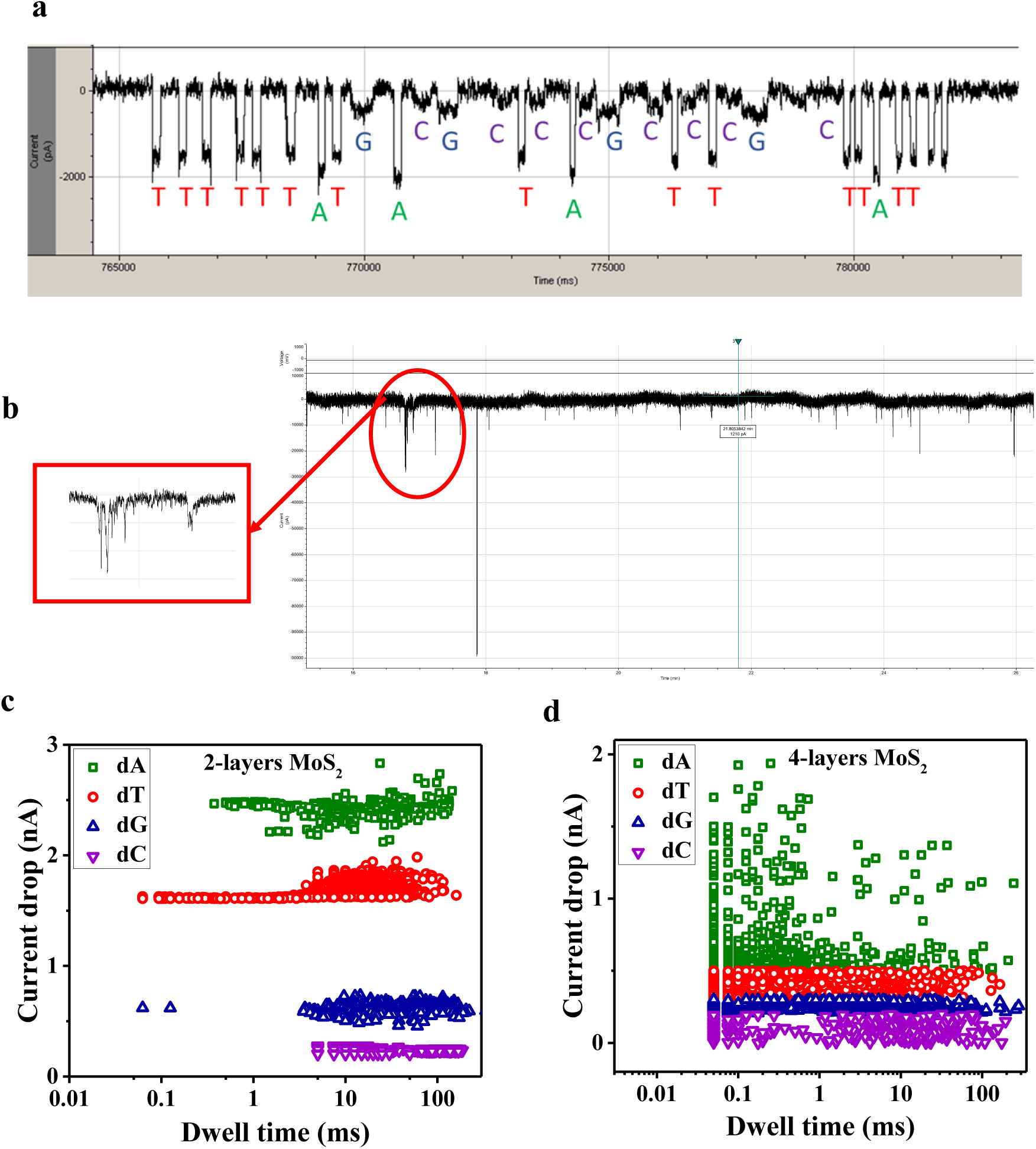
**a.** Characteristic current blockades for individual nucleotides obtained from ssDNA translocation through 2.5 nm diameter pore on 2-layers thick MoS_2_, **b.** A typical truncated ssDNA translocation profile through 2.8 nm diameter 4-layers thick MoS_2_ pore showing single nucleotide and multiple nucleotide detection peaks, **c.** Current drop vs dwell time plot for >3000 base (60 min) translocation through 2-layers thick MoS_2_: showing distinct current levels of 2.4±0.08 nA, 1.67±0.06 nA, 0.62±0.05 nA and 0.24±0.03 nA for dA, dT, dG and dC respectively with a signal-to-noise ratio> 11 (comparable to ^[14, 19, 47–48]^) and **d.** Current drop vs dwell time plot for >3000 base translocation through 4-layers thick MoS_2_: showing huge overlapping of current levels for different nucleotides; blockade currents obtained being 0.86 ± 0.33 nA for dA, 0.5 ± 0.09 nA for dT, 0.29 ± 0.07 nA for dG and 0.1 ± 0.06 nA for dC (dA: Adenine, dT: Thymine, dG: Guanine, dC: Cytosine).

### Detection rate and sensing efficiency

Table 2 shows dwell times demonstrated by 2- and 4-layers thick MoS_2_ nanopores for 60 min translocation, as extracted from Figure 3.c-d. 2-layers thick MoS_2_ successfully retards the translocation to ~67 nucleotides per minute, resolving single-nucleotide blockade signal with dwell times in the range of 0.05-140 ms. 4-layers thick MoS_2_ demonstrates dwell time, twice as that shown by 2-layers thick MoS_2_ pore, as mentioned in Table 2. Dwell times and current drops obtained for both 2-layers and 4-layers thick MoS_2_ are higher than dwell times reported for 1-layer thick MoS_2_ ^[14, 44]^.

**Table 2:** Simulated and Experimental dwell time for 4-layers and 2-layers thick MoS_2_ nanopore sensing. It can be clearly observed that there is a discrepancy in the simulated and experimental dwell time, and this can be due to the simulation parameters chosen.

| Techniques | Dwell time for 4-layers thick MoS<sub>2</sub> pore | Dwell time for 2-layers thick MoS<sub>2</sub> pore |
| --- | --- | --- |
| Simulation | 0.4-0.7 ms | 0.19-0.49 ms |
| Experiment | 0.04-350 ms | 0.05-140 ms |

Few discrepancies are observed in dwell times (see Table 2) between the simulation and experimental studies. These are due to the restrictions imposed on the nanopore system during simulation which are not strictly followed during practical experimental scenario. For example, blockade conductance in simulations depend only upon surface charge interaction between pore walls, membrane surface and molecule and on the imposed bias. Also, only vertical translation of the molecule is being considered in this case taking care of the aspect ratio and meshing issues in COMSOL Multiphysics. In experiments the blockade conductance is also influenced by the molecular conformations inside the pore, amplification factor of the tool (amplifier), noise filters, etc, which create the difference. Moreover, simulations are carried out from a distance of 4 nm away from the pore mouth for a cell size of 100 nm × 100 nm, which differs from the actual molecular trajectory and cell size (~ mm). Hence, simulations neglect the charge and kinetic interactions for a large distance in and out of the pore, giving rise to difference in results (like dwell time values).

A customized algorithm (using MATLAB R2019b) was then employed to analyse the experimental data and predict the single nucleotide sequence obtained for 2-layers thick MoS_2_ pores. The sensed nucleotide sequence was then compared with the expected sequence (of the DNA oligo sample) to recognize the number and position of nucleotides not detected. The sensing efficiency was then evaluated from the fraction of the analyte DNA strands sensed. The algorithm analyses the sensing accuracy, detection rate, nucleotide specific mean current and current deviation. The predicted sequence compared with expected oligo sequence are displayed in Figure 4.a. The results clearly show that the analyte DNA is sequenced with high accuracy, as calculated from the few undetected nucleotides obtained (indicated in yellow in Figure 4.a). Thus, based on this on this the single nucleotide sensing accuracy through 2-layers thick MoS_2_ nanopore for 3000 bases was therefore estimated by MATLAB to be about 90.22%. Figure 4.b demonstrates both ionic current and dwell time-based determination and differentiation of individual DNA bases through 2-layers thick MoS_2_ pores. A 6-33 % current deviation is observed for 4-layers thick MoS_2_ pores, whereas a 2-11 % current deviation is observed for 2-layers thick MoS_2_ asserting a more efficient single nucleotide detection through 2-layers thick pores. (see Figure 5).

**Figure 4.**
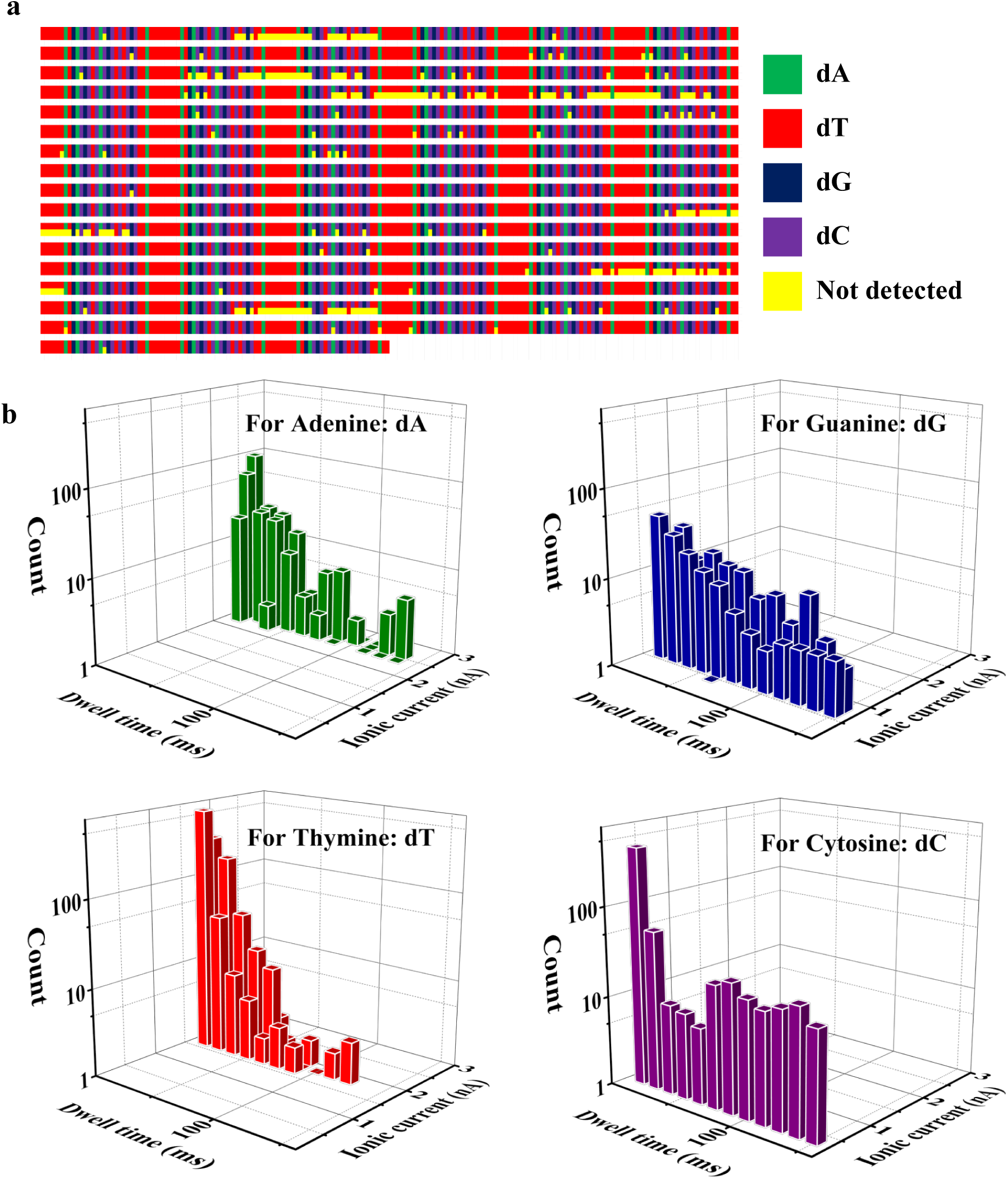
**a.** DNA sequence sensed through 2-layers thick MoS_2_ pore showing detected nucleotides: dA (green), dT (red), dG (blue) and dC (purple) and undetected nucleotides (yellow) as compared with the expected sequence, visually representing the process efficiency. It can clearly be seen from the sequence that we have very few nucleotides which were not detected. **b.** 3D histograms of sensed nucleotides with respect to ionic current and dwell times for dA, dT, dG and dC. It can be observed that four DNA nucleotides can be distinctly detected with respect to ionic current; however minor dwell time overlapping is observed which may be due to low chemical specificity of solid-state nanopore sensing.

**Figure 5.**
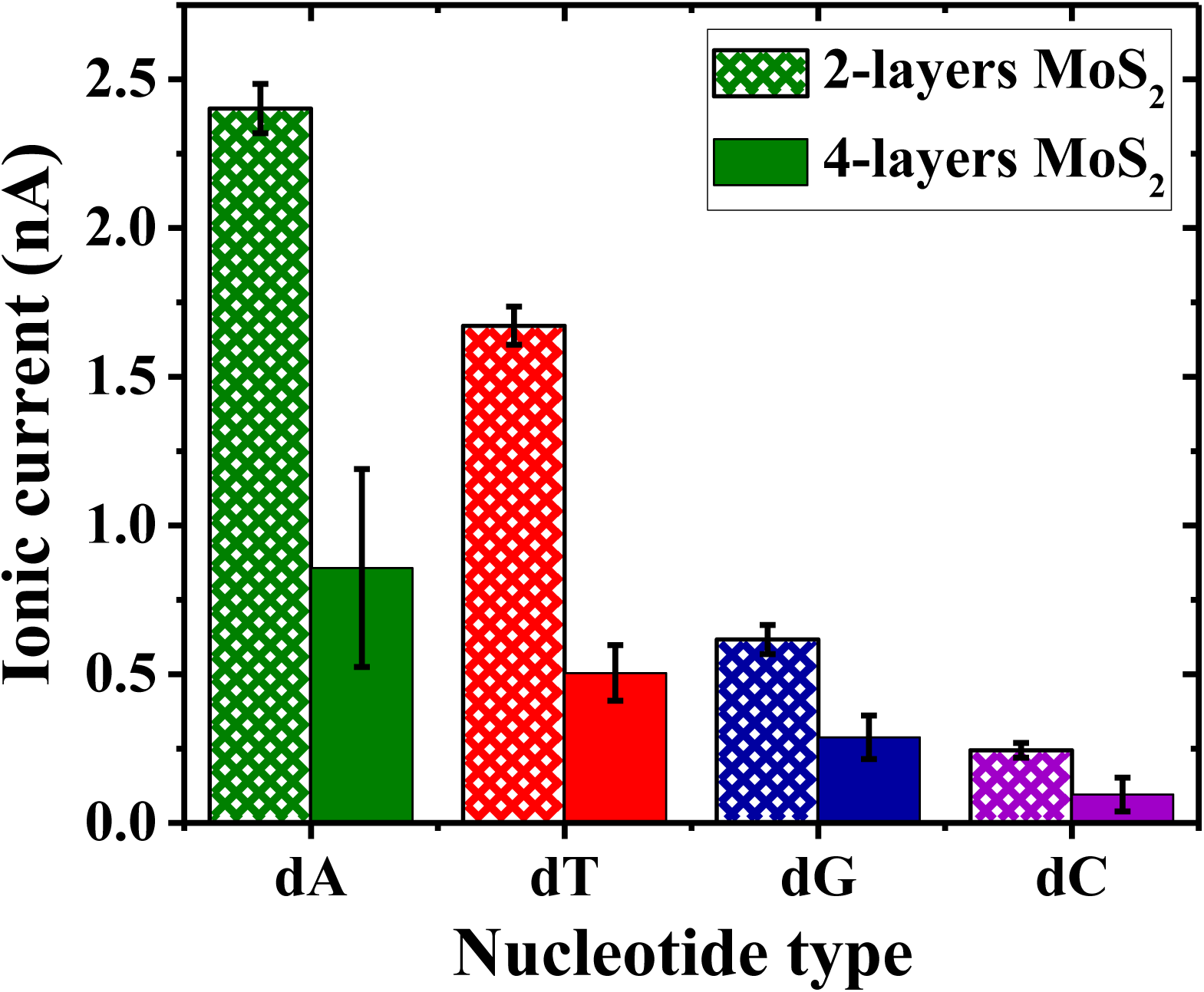
Ionic current signal magnitudes and deviation for different DNA nucleotides showing high signal accuracy displayed by 2-layers as compared to 4-layers thick MoS_2_ pores. The standard deviation (6-33 % for 4-layers thick MoS_2_ pores and 2-11 % for 2-layers thick MoS_2_ pores) reported here is from measurements on 7 pores.

The higher signal deviation for 4-layers thick MoS_2_ may be contributed to simultaneous detection of multiple nucleotides due to increased membrane thickness. The influence of surrounding molecules alters the specific signals by shifting baseline and also changing resultant charge interactions. The results summarize that interlayer potential gradient, suitable thickness compared to nucleotide dimension, good ionic current and dwell-time resolution makes 2-layers thick MoS_2_ a good choice for label-free unfunctionalized solid-state sensing of single DNA bases at picomolar concentration with high sensitivity.

Few studies have been performed in determining nucleotide type and order in DNA strand containing mixed bases by using an unfunctionalized solid-state nanopore ^[15–19]^. Read accuracy obtained in our work is similar to what is obtained by Atas et al. ^[48]^ and McNally et al. ^[49]^ Wanunu et al. (2010) ^[47]^ demonstrated miRNA sequencing at 1 fmol µl-1 (1 nM) analyte concentration, by exploiting signal resolution improvement offered by small and thin nanopores (3 nm diameter pores on 7 nm thick SiN_x_ membranes). Wanunu et al. (2010) ^[47]^ obtained 93% accuracy for RNA nucleobase sensing using complementary strand hybridized to the analyte strand. Thus, from the table it can clearly be seen that our 2-layers thick MoS_2_ nanopores with a diameter less than 3nm demonstrates high sensitivity (comparable with that obtained in literature-see Table 3) at low DNA concentrations of 40 pM even without the use of any analyte tags or additional biochemical functionalization.

**Table 3:**
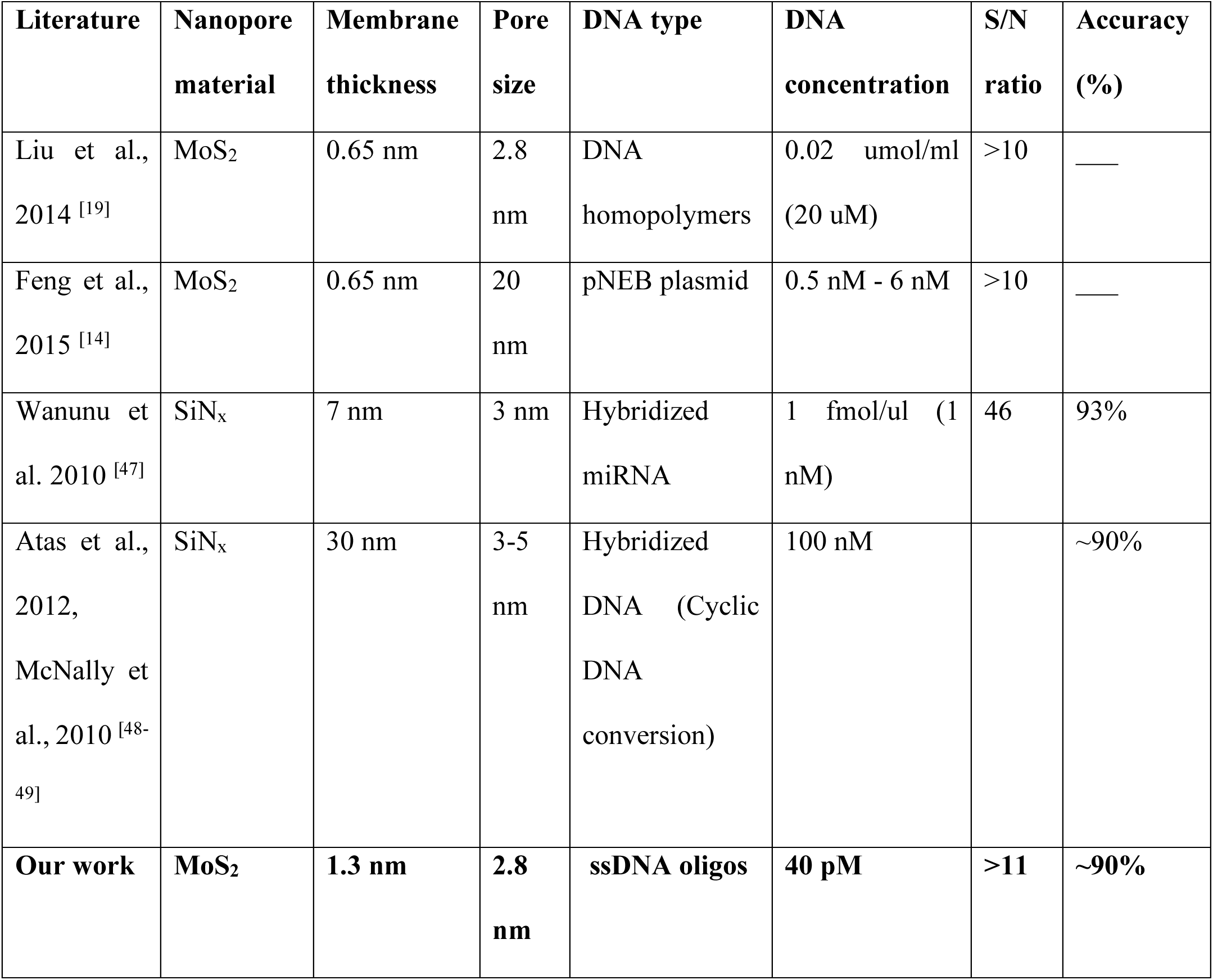
Summary of nanopore sequencing results using solid-state SiN_x_ and MoS_2_ nanopores from literature along with our results. The table summarizes the material of the pore, thickness and size and the DNA type and concentrations used along with the signal to noise ratio and accuracy of nucleotide detection.

| **Literature** | **Nanopore material** | **Membrane thickness** | **Pore size** | **DNA type** | **DNA concentration** | **S/N ratio** | **Accuracy (%)** |
| --- | --- | --- | --- | --- | --- | --- | --- |
| Liu et al., 2014 <sup>[19]</sup> | MoS<sub>2</sub> | 0.65 nm | 2.8 nm | DNA homopolymers | 0.02 μmol/ml (20 μM) | >10 | — |
| Feng et al., 2015 <sup>[14]</sup> | MoS<sub>2</sub> | 0.65 nm | 20 nm | pNEB plasmid | 0.5 nM - 6 nM | >10 | — |
| Wanunu et al. 2010 <sup>[47]</sup> | SiN<sub>x</sub> | 7 nm | 3 nm | Hybridized miRNA | 1 fmol/μl (1 nM) | 46 | 93% |
| Atas et al., 2012, McNally et al., 2010 <sup>[48-49]</sup> | SiN<sub>x</sub> | 30 nm | 3-5 nm | Hybridized DNA (Cyclic DNA conversion) | 100 nM |  | ~90% |
| **Our work** | **MoS<sub>2</sub>** | **1.3 nm** | **2.8 nm** | **ssDNA oligos** | **40 pM** | **>11** | **~90%** |

## Experimental detail

### Fabrication of MoS_2_ nanopores

For fabricating SiN_x_ membranes double-sided LPCVD (low stress) silicon nitride coated 4 inch, 525 ± 25 µm, <100> silicon wafers were purchased from Rogue Valley Microdevices. For MoS_2_ membranes low defect density highly oriented 2H-phase single crystal was used which was procured from 2D semiconductors inc. Single-stranded DNA oligos with 25.5 n-moles (of a cutomized nucleotide sequence without any end functionalization) containing 30 mixed nucleotides in 250 µl solution was procured from idtdna.com. All of the other reagents were obtained from Fisher Scientific. Solid-state free-standing SiN_x_ membrane was fabricated using microfabrication techniques like lithography and etching (dry and wet). The MoS_2_ nanopores were fabricated by starting with the free standing SiN_x_ membrane as substrate. A 200 nm diameter hole was patterned at the center of the membrane by EBL followed by Reactive Ion Etching (RIE). Then, 2-layers (~ 1.3 nm) and 4-layers (~ 2.5 nm) thick MoS_2_ flakes were mechanically exfoliated (scotch-tape) on the 200 nm diameter hole to form the free standing MoS_2_ membrane using a dual stage microscope for better alignment and centering. Size of MoS_2_ flakes were found to be around 20 µm repeatedly. Then the nanopore was drilled on top by JEOL JEM-ARM200CF Scanning/ Transmission Electron Microscope (S/TEM). We were able to obtain pores in the size range of 2- 4 nm diameter repeatedly based on our application requirement. Vigorous imaging on the same pore is avoided to prevent pore expansion due to prolonged beam exposure. Maintaining a good control over fabrication parameters we could achieve pores with an error of about 1-2 A^0^ in diameter, every time. The steps of fabrication for both membrane materials are given in **Figure S5.a** (see **Figure S6** for the experimental images of fabrication steps).

### Reagent preparation

300 mM filtered KCl solution was prepared and buffered with 3 mM Tris-HCl was prepared (pH=8) and stored (to be used for measurement of ionic conductance). The latter solution was then diluted by DI water to 100 mM for picomolar concentration analyte translocation experiments. The stored KCl solution was degassed for 90 minutes in vacuum. Before experiments, the temperature of degassed KCl was ensured to be at room temperature. Analyte solutions at 40 pM concentration was prepared by repeated dilution of 8 µl of purchased DNA solution.

### Custom-designed cell assembly and cleaning

Half cells made of Poly-tetra-fluoro-ethylene (Teflon) were used as electrolytic chambers. Teflon having a good electro-chemical resistance and hydrophobicity helps maintain clean and insulated environment, required for accurate sensing. Polydimethylsiloxane (PDMS) gaskets were used to seal the half cells in order to prevent cross flow of electrolyte in between cells other than through nanopore. PDMS gaskets reduce capacitive noise during ionic current measurements. Sonication-assisted thorough cleaning of half cells was done before each experiment to remove any KCl residue. Bias across the membrane was applied using Ag/AgCl electrodes dipped in the electrolyte contained in the half cells. Electrodes were functionalized with chloride by treating them with ethanol, DI water and dipping them in bleach for about 1 hour. Electric interference was prevented by isolating the entire assembly in a Faraday cage purchased from Warner instruments. For ionic current measurements both the cell chambers were filled with 0.3 M KCl solution. **Figure S5.b** of Supplementary information shows the experimental setup used for all experiments. Single channel recordings were obtained to characterize the solid-state nanopores. The ionic blockades induced by translocating DNA nucleotides were filtered by 2kHz 8-pole Bessel filter, amplified by Axopatch 700B and finally digitized by Digidata 1550B.

Ionic current through the nanopore was measured at voltage swept from −200 mV to 200 mV to measure the ionic conductance. Each pore was measured 7 times to obtain a stable conductance after repeated cleaning and the average conductance starting from the third measurement was reported.

### Pore Cleaning and mounting

The membranes are cleaned three times by pulling acetone under vacuum for 30 minutes each, with the acetone solution being replaced by a new solution every time to avoid drying up of acetone at the nanopore. Then the same process is repeated four times using IPA and once using ethanol, each cycle was carried out for 20 minutes. During initial measurements, 4 cycles of IPA clean was found to be optimum for removing all acetone and achieve a low noise level by avoiding residues.

Before experiments, the nanopore immersed in stored degassed KCl was electrically conditioned at constant 100 mV voltage till the noise level is reduced to an accepted minimum.

## Conclusion

In this study, we have conducted detailed simulation and experimental work to understand the behaviour of different number of layers of 2D material like MoS_2_ for its application in nanopore fabrication and sensing. Both our simulation and experimental studies clearly indicate that the multilayer thick MoS_2_ nanopore shows better sensitivity than a single layer one. It is clearly observed that even number of layers as the interlayer coupling potential provides increased electrophoretic pull. We observe that the 2-layers thick MoS_2_ nanopore displays greater single nucleotide detection accuracy of ~ 90% with signal to noise ratio greater than 11 at 40 pM DNA concentrations compared to 4-layers. Thus, we have conclusively demonstrated a high accuracy and signal to noise ratio for 2-layers thick MoS_2_ nanopores with diameters less than 3 nm without any functionalization for analyte concentrations as low as 40 pM. It can thus be concluded that 2D materials like MoS_2_ can be tuned to obtain high quality nanopores for ultra-sensitivity and accuracy which can be utilised effectively for nanopore applications for detection of low concentration analytes.

## Notes

### Competing Interest Statement

The authors have declared no competing interest.

